# NFIA regulates granule recruitment and exocytosis in the adult pancreas

**DOI:** 10.1101/2019.12.24.885020

**Authors:** Marissa A. Scavuzzo, Jolanta Chmielowiec, Jessica Teaw, Diane Yang, Matthew C. Hill, Andrea R. Waksmunski, Lita Duraine, Joan Camunas-Soler, Xiaoqing Dai, Jordon C. King, Stephen R Quake, Patrick E. MacDonald, Andre Catic, Malgorzata Borowiak

## Abstract

After food ingestion, pancreatic cells secrete zymogen and hormone-containing granules to precisely control digestion and blood glucose levels. Identifying regulators of this process is paramount to combatting multiple pancreatic diseases. Here we show that pancreatic deletion of the transcription factor nuclear factor IA (NFIA) leads to hyperglycemia, hypoinsulinemia, and hypolipidemia. Surprisingly, insulin and digestive enzymes are produced in the absence of NFIA, however, they are not secreted properly and instead accumulate inside pancreatic cells. In NFIA-deficient mice we saw a reduction of insulin granules in the ready releasable pool and the first-phase insulin response was impaired. We found that NFIA binds to and activates *Rab39b,* a Rab GTPase critical for exocytosis. Re-expression of *Rab39b* in NFIA knockout islets restored glucose-stimulated insulin secretion. In sum, the NFIA-Rab39b axis regulates pancreatic physiology through <u>granule recruitment and docking</u>, linking NFIA to a new process with potential effects in diabetes, pancreatitis, and lipid disorders.

## Introduction

The careful choreography of vesicle trafficking proteins ensures that secretory vesicles are transported to the membrane for exocytosis upon stimulation. This process includes the coordination of different Rab GTPases, which can act alone or in concert with other proteins to chauffer vesicles to specific locations within the cell. This process is of utmost importance in the pancreas, a dedicated secretory organ in which different cell types calibrate precise amounts of zymogen and hormone-containing granules to exocytose for regulation of digestion and blood glucose levels.

In response to elevated blood glucose levels, endocrine beta cells secrete insulin, which binds to receptors on peripheral tissues to induce glucose uptake (Sonksen and Sonksen, 2000). Insulin is the only hormone capable of reducing blood glucose levels and is solely produced by pancreatic beta cells. Defects in beta cell function leading to inadequate insulin production or secretion can lead to diabetes (Weir and Bonner-Weir, 2004). Consequently, the mechanisms orchestrating insulin secretion from beta cells must be tightly regulated to keep blood glucose levels at homeostasis.

Acinar cells surround endocrine islets and constitute the majority of the pancreas, secreting more proteins than any other cell type in the body. Upon feeding, acinar cells release digestive enzymes stored as inactive zymogens (Case, 1978). Disruption to zymogen secretion can lead to pancreatitis, with the pancreas becoming inflamed, necrotic, and fibrotic, which in turn affects digestion and can lead to pancreatic cancer (Krah and Murtaugh, 2016). Therefore, the identification and characterization of genes involved in regulating insulin and zymogen granule secretion is essential to understand and treat a multitude of pancreatic diseases.

Insulin is produced and packaged into secretory granules, followed by the translocation of approximately 5% of these granules from a reserve pool to dock at the plasma membrane ready for release (the “ready releasable pool” of granules) (Hao et al., 2005). Rising levels of blood glucose are detected by uptake of glucose through the glucose transporter (Glut2 in mice) before conversion to ATP and other metabolites (Fu et al., 2013). The resulting rise in ATP inhibits ATP-sensitive potassium channels, causing depolarization of the membrane and opening of voltage-gated calcium channels. Depolarization causes what is described as the first-phase insulin response, consisting of the immediate exocytosis of membrane-docked granules (Hao et al., 2005). A second amplification phase follows, in which granules stored in the reserve pool are readied for release. Transient association with protein interacting with C-kinase 1 (Pick1) leads to the maturation and recruitment of granules to the membrane for secretion (Cao et al., 2013). Pick1 must interact with Rab GTPase Rab39b in order for AMPA receptors to traffic to the membrane, however what regulates the association of Pick1 with granules in the pancreas is unknown (Mignogna et al., 2015).

We previously described the transcription factor nuclear factor I-A (NFIA) to regulate cell fate determination during pancreatic development (Scavuzzo et al., 2018), however the adult functions of NFIA have yet to be defined. Genome wide association studies illustrate that mutations in *NFIA* are associated with diabetes mellitus and hyperlipidemia (Dupuis et al., 2010; Lee et al., 2013; Nettleton et al., 2010; Teslovich et al., 2010), leading us to hypothesize that NFIA plays a role in the adult pancreas to regulate pancreatic physiology. Here, we selectively deleted NFIA from pancreatic cells in mice, which led to hyperglycemia, hypoinsulinemia, and hypolipidemia. Surprisingly, insulin and digestive enzymes like amylase were produced, however, their secretion was diminished. Instead, we found that enzymes accumulated inside acinar and beta cells in the pancreas. We found that NFIA-deficient beta cells harbor a trafficking defect with impaired recruitment and docking of insulin granules to the plasma membrane, with a reduction in the ready releasable pool and defects in the first-phase insulin response. By overlapping CUT&RUN and RNA-sequencing data, we pinpointed genes NFIA directly targets, including *Rab39b,* which coats granules with Pick1 for regulated exocytosis. We validated this finding by demonstrating that Rab39b is decreased and Pick1 is mis-localized in NFIA-knockout beta cells. We further showed that re-expression of *Rab39b* in NFIA knockout islets restores insulin secretion after glucose stimulation. This work identifies unique functions of the developmental transcription factor NFIA in regulating exocytosis to control adult pancreatic physiology.

## Results

### Pancreatic deletion of the transcription factor NFIA to determine its role of in diabetes and digestive abnormalities

Diabetes mellitus and lipid disorders are highly hereditary (Ali, 2013; Shapiro, 2000), thus the identification of genetic variants in patients can inform downstream genetic studies and lead to a mechanistic understanding of disease etiology. There is evidence of associations between genetic variants in *NFIA* and diabetes and elevated high-density lipoprotein (HDL) cholesterol levels in genome-wide association studies (GWAS) (**Figure 1A-B, S1A;** (Dupuis et al., 2010; Hu et al., 2015; Lee et al., 2013; Nettleton et al., 2010; Teslovich et al., 2010)). To determine whether *NFIA* is expressed in the adult pancreas, we immunostained the adult mouse and human pancreas for NFIA. We found *NFIA* present in all three major components of the pancreas, in endocrine, ductal, and acinar cells (**Figure 1C**). The expression of NFIA in pancreatic islets by immunohistochemistry appeared heterogeneous; to determine whether NFIA was indeed expressed in islets, we isolated mouse islets and whole pancreas. By Western blot analysis we observed high expression of NFIA in mouse islets relative to beta-actin and in comparison to total pancreas (**Figure S1B**). These findings are consistent with an increase in human *NFIA* mRNA expression from human fetal week 20 to adulthood (**Figure S1C**). Because *NFIA* germline knockout mice die at birth, we generated conditional NFIA mutant mice with lox_p_ sites flanking the second constitutive exon containing DNA binding and dimerization domains (Scavuzzo et al., 2018). After crossing to Pdx1-cre mice to delete NFIA in pancreatic cells, which we call NPKO (<u>N</u>FIA <u>P</u>ancreatic <u>K</u>nock<u>O</u>ut), we observed a shift in the proportion of cell types forming, with markedly fewer endocrine cells (**Figure 1D, S1D,** (Gu et al., 2002)). We also observed elevated fibrosis in the adult NPKO pancreas, indicating dysfunction of acinar cells (**Figure 1E**). While 12-week-old NPKO mice have no change in body weight compared to controls (**Figure S1E**), they exhibited a reduced pancreas size relative to body size (**Figure S1F, S1G**). Thus, NFIA mutations not only correlate with defects in blood glucose homeostasis and digestion in patients, but NFIA is specifically expressed in the cells that regulate blood glucose levels and digestion in the adult pancreas, suggesting a causal relationship.

**Figure 1.**
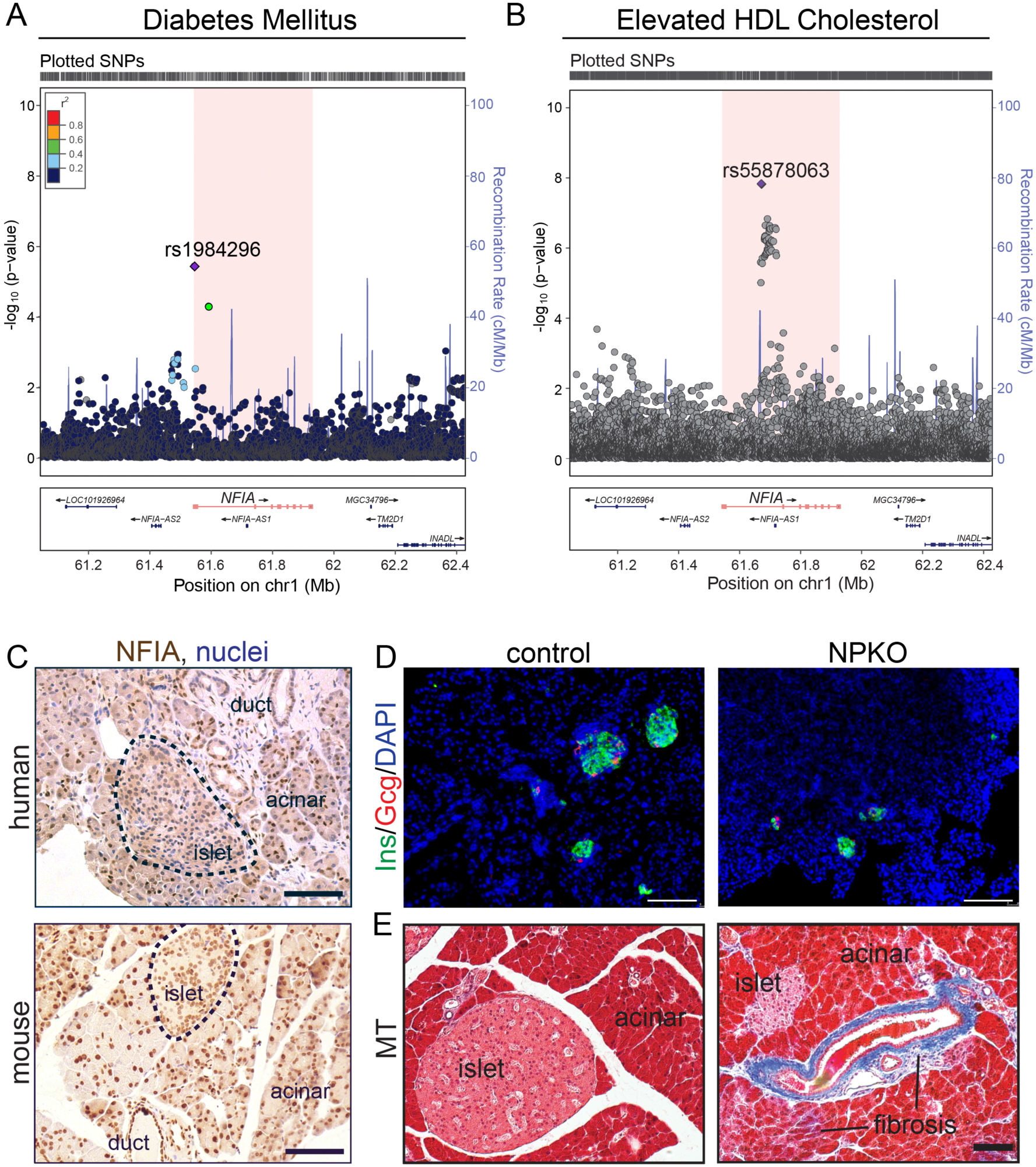
The transcription factor NFIA is associated with pancreatic diseases. A. LocusZoom plot highlighting suggestive diabetes variant (rs115855396, p = 6.19×10^-6^) upstream of *NFIA* in the National Heart, Lung and Blood Institute (NHLBI) Family Heart Study (FamHS). P-values were previously generated in a genome-wide association study of individuals taking medication for diabetes or glucose in the NHBLI FamHS at Visit 1 (NCBI dbGAP analysis accession: pha003059). rs115855396 is highlighted in purple. Linkage disequilibrium (LD) estimates (r^2^) are based on the European population from the 1000 Genomes Project (November 2014 release). B. LocusZoom plot of p-values for *NFIA* variants in GeneATLAS. P-values for variants within *NFIA* +/- 500 kilobases were downloaded from the GeneATLAS, which is a publicly accessible database of UK Biobank summary statistics from genome-wide association studies (Canela-Xandri et al. 2018). The results shown correspond to the GWAS of diabetes in GeneATLAS (UK Biobank field: 20002; Field codes: 1220, 1223, 1521, 1221, 1222), which was performed on 21,105 diabetes cases and 431,159 controls. The lead variant (rs1984296, 3.77×10^-6^) is depicted as a purple diamond. The color scheme corresponds to LD estimates (r^2^) in the European population from the 1000 Genomes Project (November 2014 release). C. Immunohistochemistry showing NFIA (in brown) expression in adult mouse (8.5w) and human adult (64-year old) pancreas in endocrine, duct, and acinar cells (with nuclear hematoxylin co-stain in blue). D. Immunofluorescent staining of d20 control and NPKO pancreas showing beta cells (Ins in green) with alpha cells (Gcg in red). Nuclei are marked by DAPI in blue. Scale bars=100 μm. E. Gross morphology of adult control and NPKO pancreas by Masson’s trichrome (MT) histological staining. Scale bars=75 μm.

### NFIA-deficient mice develop hyperglycemia and digestive defects

To determine whether NFIA functions in adult beta or acinar cells, we physiologically assessed NPKO mice for defects in blood glucose homeostasis and digestion. Adult NPKO mice fed *ad libitum* developed elevated blood glucose levels compared to control mice; 82% of mice displayed hyperglycemia with blood glucose levels over 150 mg/dL (**Figure 2A**). After 7 hours of fasting, control mice exhibited a significant drop in blood glucose levels, while the hyperglycemic state of NFIA-deficient mice persisted even after fasting (94% elevated, **Figure 2A**). Blood glucose levels of NPKO mice continued to be elevated compared to controls after fasting for 16 hours (**Figure 2A**). We next assessed the ability of NPKO mice to clear glucose from the blood after fasting and glucose challenge. The NPKO mice were glucose-intolerant compared to control mice, with raised blood glucose levels throughout the glucose tolerance test (**Figure 2B,C**). Controlled fasting and re-feeding further illustrated the physiological defects in NPKO mice, with elevated blood glucose levels in both states (**Figure 2D,E; Figure S2A**).

**Figure 2.**
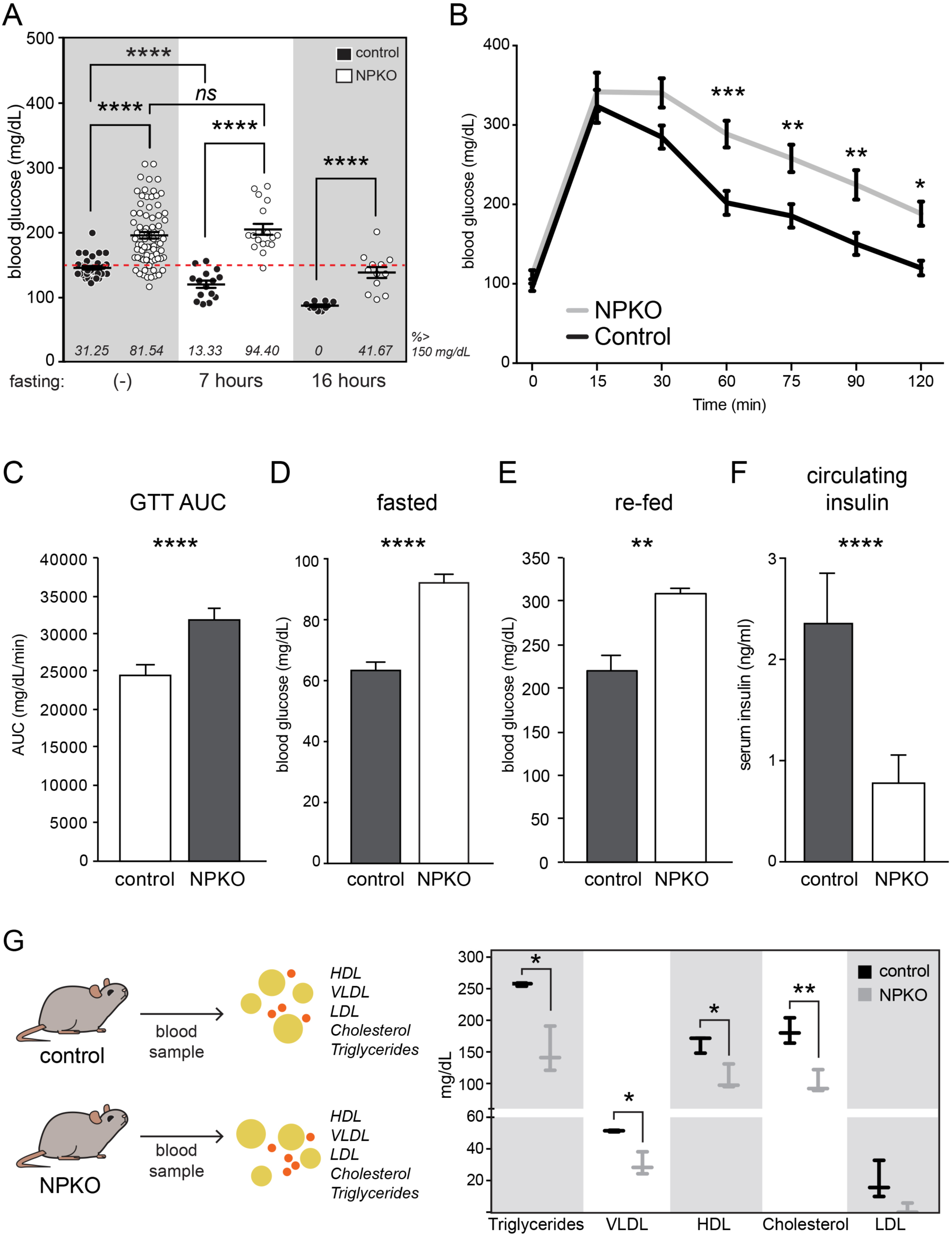
Conditional deletion of pancreatic NFIA leads to hyperglycemia, hypoinsulinemia and hypolipidemia. A. Blood glucose concentrations, measured in randomly fed (-), short-term fasted (7 hours), and long-term fasted (16 hours) adult NPKO mice compared to age and gender matched controls. Black dots indicate control mice while white dots indicate NPKO mice. Each symbol represents one mouse (no fasting: N=32 control mice, N=78 NPKO; 7 hours fasting: N=16 control mice, N=19 NPKO mice; 16 hours fasting: N=10 control mice, N=12 NPKO mice). Red dashed line indicates 150 mg/dL, the cutoff for normoglycemia in mice. The percentage of mice above the cutoff for normoglycemia is shown below each measurement. Error bars show SEM, \*\*\*\**p<*0.001; *ns=*not significant between mice. B. Intraperitoneal glucose tolerance tests (GTTs) of adult control (shown in black) and NPKO (shown in gray) mice show elevated blood glucose in the absence of NFIA. N=15 Red dashed line indicates 150 mg/dL, the cutoff for normoglycemia in mice. N=16 control and N=15 NPKO mice. Error bars show SEM, \*\*\**p<*0.005; \*\**p<*0.01; \**p<*0.05 between mice. C. Quantification of area under the curve (AUC) for GTTs. Error bars show SEM, \*\*\*\**p<*0.001 between mice. D. Blood glucose concentrations after overnight fasting. N=5 control and N=3 NPKO mice. Error bars show SEM, \*\*\*\**p<*0.001 between groups. E. Blood glucose concentrations after re-feeding. Mice were fasted overnight, injected with glucose, and assessed 30 minute later. N=5 control and N=3 NPKO mice. Error bars show SEM, \*\**p<*0.01 between mice. F. Random blood serum insulin levels of mice fed with standard chow in adult control (shown in gray) and NPKO (shown in white) mice. NPKO mice have decreased levels of insulin in the blood stream. N=9 mice per genotype. Error bars show SEM, \*\*\*\**p<*0.001 between mice. G. Lipid levels in the blood serum of adult control (in black) and NPKO (in gray) showing mean with upper and lower bounds. N=3 mice per genotype. Error bars show SEM, \*\**p<*0.01; \**p<*0.05 between mice.

In NPKO mice, NFIA is deleted from all pancreatic cell types. To determine whether the defects in blood glucose homeostasis were primary to beta cells or rather a secondary effect, we crossed NFIA^fl/fl^ mice to *Ngn3*-cre mice to delete NFIA from all endocrine cells. We found elevated blood glucose levels in NFIA^fl/fl^; *Ngn3*-cre mice (**Figure S2B,** (Schonhoff et al., 2004)). We next crossed NFIA^fl/fl^ mice to *Ins2*-cre mice to generate NFIA beta cell knockout mice and found that hyperglycemia persisted in NFIA beta cell knockout mice, indicating an intrinsic beta cell defect in glucose homeostasis (**Figure S2C,** (Postic et al., 1999)). Pancreatic beta cells produce insulin to lower blood glucose levels. As anticipated, we found that mice without pancreatic NFIA had significantly lower insulin levels in the blood stream (**Figure 2F**), but no difference in insulin sensitivity (**Figure S2D, S2E**). Overall, NPKO mice exhibited a diabetic phenotype, with inadequate levels of circulating insulin and chronically elevated blood glucose levels.

In addition to an association with glucose homeostasis, we found that NFIA mutations in patients were associated with abnormal triglyceride, HDL cholesterol, and lipid levels. Further, we observed NFIA expression in digestive acinar cells. We found that fresh NPKO mouse feces float compared to control mouse feces, indicating an accumulation of fat in the stool (**Figure S2F**). These observations suggested a disruption to digestion in NPKO mice, with impediment to lipid breakdown. Indeed, lipid panel analysis of blood serum showed a significantly decreased level of circulating triglycerides, very low-density lipoprotein (VLDL), HDL, and cholesterol in NPKO mice compared to controls (**Figure 2G**), suggesting that NPKO mice cannot efficiently digest and breakdown lipids, so they are instead retained in the stool. These results indicate that pancreatic NFIA may also regulate digestion.

### Loss of pancreatic NFIA leads to granule accumulation and reduction of the insulin ready releasable pool

To address whether the hypolipidemia, hypoinsulinemia, and hyperglycemia in NPKO occurred due to a decrease in zymogen or insulin production or secretion, we next assessed the content of granules within the pancreas. Transmission electron microscopy (TEM) revealed a remarkable accumulation of zymogen granules within acinar cells in the NPKO pancreas (**Figure 3A, 3B**). We observed a higher number of large granules (**Figure 3C, S3A**), which were spread throughout acinar cells rather than polarized towards the apical membrane (**Figure 3D**), suggesting increased maturation or lifespan of zymogen granules in NPKO mice. While we found a slight decrease in amylase activity in the blood serum of NPKO mice (**Figure S3B**), we found a significant increase in amylase activity within the pancreas itself (**Figure 3E**), showing that amylase accumulated within acinar cells instead of entering the blood stream.

**Figure 3.**
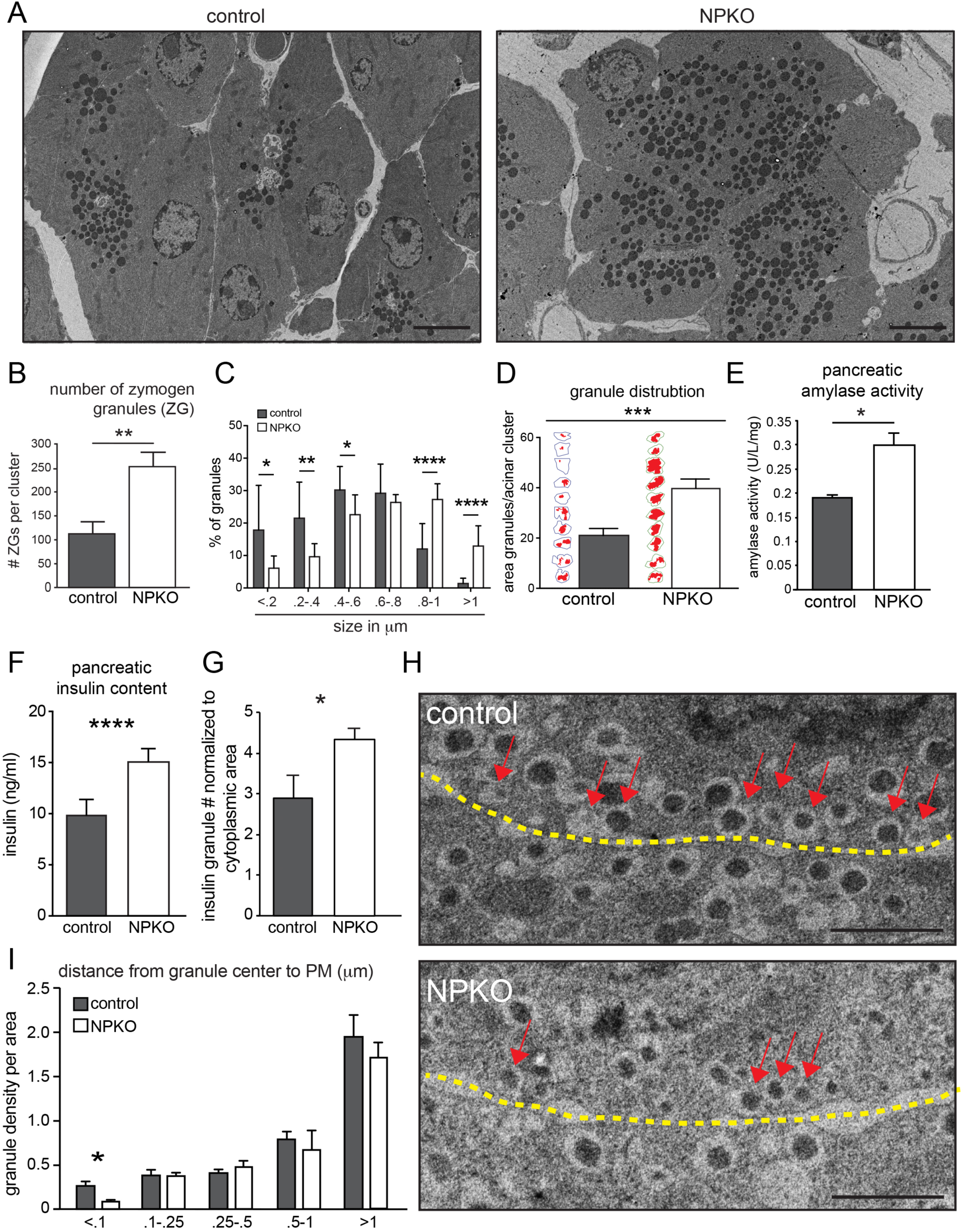
Zymogen and insulin granule trafficking is impaired in NFIA-deficient acinar and beta cells. A. Transmission electron micrograph of pancreatic acinar cells in adult control and NPKO mice. B. Increased number of zymogen granules per acinar cluster in adult NPKO mice, N=2 mice per genotype. Error bars show SEM, \*\**p<*0.01 between cells. C. Increased zymogen granule size in area (μm^2^) in adult NPKO mouse pancreatic acinar cells. N=4 control and N=2 NPKO mice, n=893 control granules and n=835 granules in NPKO. Error bars show SEM, \*\*\*\**p<*0.001, \*\**p<*0.01, \**p*<0.05 between cells. D. Increased zymogen granule distribution throughout the space of the acinar cluster in adult NPKO mice, as measured by the area of the granules normalized to the area of the acinar cluster. Distribution maps shown normalized to acinar cluster area. N=4 control and N=2 NPKO mice. Error bars show SEM, \*\*\**p*<0.005 between cells. E. Increased amylase activity in the pancreas of NPKO and control mice. N=3. Error bars show SEM, \**p<*0.05 between mice. F. ELISA measurements of insulin content in the pancreas of adult NPKO compared to control mice. NPKO mice accumulate insulin in the pancreas, N=10 mice per genotype. Error bars show SEM, *\*\*\*\*p*<0.001 between mice. G. Total number of insulin granules in control and NPKO mice beta cells, normalized to beta cell cytoplasmic area. NPKO mice exhibit an accumulation of insulin granules, N=3 control and N=2 NPKO mice. Error bars show SEM, \**p<*0.05 between mice. H. Transmission electron micrograph of beta cells in adult control and NPKO mice. Dotted yellow line represents the plasma membrane. Red arrows show docked insulin granules at the plasma membrane. Scale bar=500 nm. I. The distance from the plasma membrane to the center of insulin granules binned into 5 groups (0-100, 100-250, 250-500, 500-1000, greater than 1000 nm insulin granules). NPKO mice have fewer insulin granules in the docked granule zone (0-100 nm) compared to control mice. Data is shown as percentage of granule density (granules/binned area) relative to cytoplasmic area (area of the beta cell cytoplasm), N=3 control and N=2 NPKO mice. Error bars show SEM, *\*p<*0.05 between mice.

To functionally assess the consequence of zymogen granule accrual in NPKO acinar cells, we challenged control and NPKO mice with repeated caerulein injections prior to collecting blood to assess amylase activity levels. Caerulein, which mimics the action of cholecystokinin, is a secretagogue that at high or repeated doses causes hyper-secretion and premature activation of zymogen granules inside acinar cells. Premature activation of zymogens within the pancreas can lead to pancreatitis (Thrower et al., 2006), and we observed elevated fibrosis in the NPKO pancreas (**Figure 1E**). Following five intraperitoneal injections of caerulein, we found that amylase activity in NPKO mouse blood serum increased over time relative to control mice, along with markedly increased inflammation in the pancreas suggesting a propensity to form pancreatitis (**Figure S3C-D**). This increase at late timepoints and massive inflammatory response likely reflects the release of granules into the circulation of NPKO mice after acinar cell death.

Overall, we found that the number and size of zymogen granules increased in NPKO pancreatic acinar cells and that they were mis-localized, while amylase levels increased within the pancreas but not in circulation. Secretogogue-induced pancreatitis increased amylase levels in the blood serum of NPKO mice at late stages compared to controls, indicating a defect in zymogen granule trafficking and secretion.

NPKO mice developed diabetic phenotypes with significantly elevated blood glucose levels and decreased circulating insulin (**Figure 2**). However, we found that NPKO mice had an increased amount of insulin within the pancreas (**Figure 3F**). These results suggest that insulin is trapped inside the pancreas of NPKO mice, unable to be trafficked or secreted and instead accumulating within beta cells. By TEM, we found an increase in the total number of insulin granules in NFIA-deficient beta cells, further supporting the improper release of these granules (**Figure 3G**).

Insulin granules in pancreatic beta cells are localized either in a ready releasable pool at the membrane or in a reserve pool (Bratanova-Tochkova et al., 2002) (**Figure S4A**). We saw a 30% reduction in the number of granules docked at the plasma membrane, which concurs with the diminished response to glucose stimulation (**Figure 3H**). This reduction of insulin granule docking in NFIA-deficient beta cells was specific to the ready-releasable pool, as insulin granules further from the plasma membrane were equivalent in density to controls but showed a marked reduction within 100 μm of the plasma membrane (**Figure 3I**). By TEM, we saw no change in the proportion of dense insulin granules in NPKO mice (**Figure S3E**), suggesting that NFIA regulates granule recruitment or docking. Together, loss of NFIA in pancreatic acinar and beta cells leads to accumulation of granules within the cells and higher levels of enzymes and hormones within the pancreas but less secreted into the bloodstream.

*NPKO beta cells are dysfunctional with an impaired first phase insulin response* Pancreatic deletion of NFIA causes a decrease in the formation of endocrine cells ((Scavuzzo et al., 2018), **Figure 1F**). Thus, the hyperglycemia in NPKO mice could be caused by reduced beta cell number or by impaired beta cell function. As we found elevated blood glucose levels in mice with NFIA deleted only in beta cells after cell fate specification, we sought to determine the level of insulin secretion per beta cell to evaluate whether this phenotype was related to cell number, structural defects, or cellular function. Therefore, we isolated whole islets from control and NPKO mice and performed glucose stimulated insulin secretion assays (GSIS) before islet dissociation, insulin ELISA, and normalization of insulin levels to cell number (**Figure 4A**). Loss of NFIA caused a defect in insulin secretion after both glucose and KCl stimulus, indicating that these beta cells are unable to properly exocytose insulin (**Figure 4B, S4B**).

**Figure 4.**
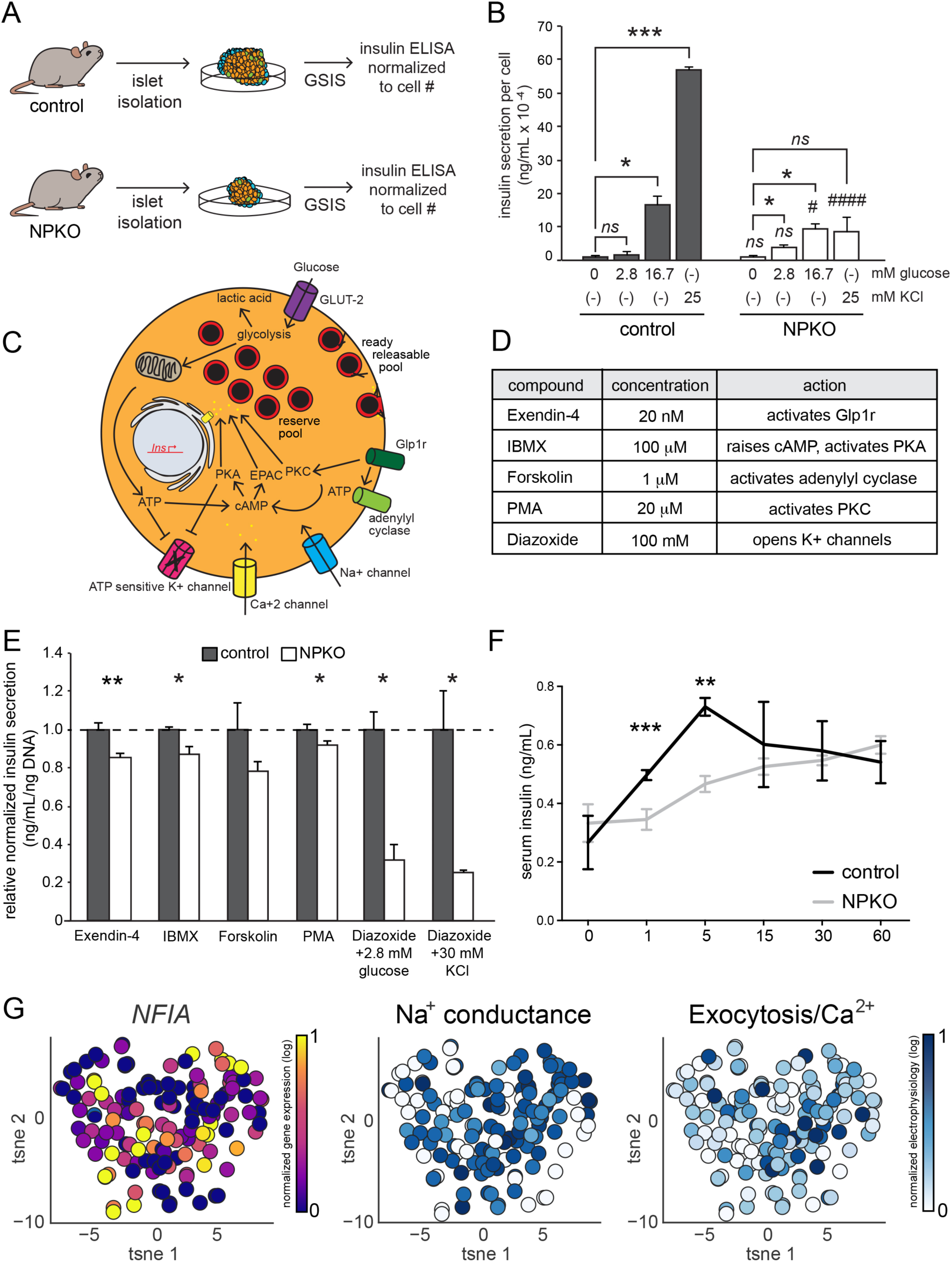
Loss of NFIA leads to defects in beta cell physiology. A. Schematic shows assessment of beta cell dynamics by isolating islets from adult control and NPKO mice and performing glucose stimulated insulin secretion (GSIS) assays normalized to cell number. B. Insulin secretion from adult control and NPKO beta cells after basal (2.8 mM), high (16.7 mM) glucose, and KCl (25 mM) stimulus. NPKO exhibit deficiencies in insulin secretion in response to stimuli, indicating a secretory defect. N=8 mice per genotype, n=2 technical replicates for insulin ELISA. Error bars show SEM compared between mice, \*\*\**p<*0.005, \**p*<0.05, ####*p*<0.001 between control and NPKO 25 mM KCl stimulation, #*p*<0.05 between control and NPKO 16.7 mM glucose stimulation, *ns=*not significant. C. Schematic of a beta cell showing mechanisms of insulin granule exocytosis. D. Compounds used in secretagogue assay and corresponding concentrations and mechanisms of action. E. Secretagogue assay shows diminished response of NPKO islets regardless of stimulation route. N=4 mice per genotype. Error bars show SEM, \*\**p<*0.01, \**p*<0.05 between mice. F. Serum insulin levels were measured to test first-phase insulin secretion after 16 hours fasting and glucose stimulus in control (shown in black) and NPKO (shown in gray) mice. NPKO mice exhibit a diabetic phenotype with limited first-phase response to glucose. N=3 mice per genotype, n=2 technical replicates for insulin ELISA. Error bars show SEM, \*\*\**p<*0.005, \*\**p*<0.01 between mice. G. tSNE projection of patch-seq beta-cells from nondiabetic donors incubated at high glucose conditions (5-10 mM). Color shows scaled expression of *NFIA* (on left) and electrophysiological parameters (center and right) for each cell.

We next sought to determine whether the decrease in insulin secretion was consistent with stimulation by other secretagogues that stimulate insulin secretion through mechanisms other than glucose (**Figure 4C**). NPKO islets showed a diminished response to Exendin-4, an incretin mimetic that activates Glp1r and promotes insulin secretion through activation of the cAMP binding protein EPAC (Goke et al., 1993), 3-Isobutyl-1-methylxanthine (IBMX), which activates cAMP, protein kinase A (PKA) signaling, and calcium release from the endoplasmic reticulum (Grill et al., 1975), Forskolin, which activates adenylyl cyclase (Seamon et al., 1981), and phorbol 12-myristate-13-acetate (PMA), which activates protein kinase C (PKC) (Vandenbark et al., 1984) (**Figure 4D, 4E, S4C**). Finally, we also observed reduced insulin secretion after treatment with Diazoxide, which activates potassium channels to hyperpolarize cells (Arkhammar et al., 1987), in combination with 2.8 mM glucose or 30 mM KCl in NPKO islets compared to controls. This reduction in insulin secretion shows that activation of potassium channels worsens the phenotype caused by NFIA deletion even after co-stimulation with KCl, indicating that there may be a decrease in the NPKO beta cell ready releasable pool of insulin (**Figure 4D, 4E, S4C**).

To determine whether there were metabolic changes in NPKO beta cells, we measured the mitochondria area from TEM images, which showed no significant differences in mitochondria size or number (**Figure S4D-G**). In line with these findings, we also did not see any defects in cellular respiration, and in fact observed elevated maximal respiration in NPKO islets (**Figure S4H-I**). These results from *ex vivo* islets illustrate that the hyperglycemia in NFIA-deficient mice was not caused solely by a decrease in endocrine cell number but also due to impaired beta cell function.

If beta cells indeed have a defect in secretion after loss of NFIA, then we would expect to see a defect in the first-phase insulin response after glucose stimulation (**Figure S4A**). In order to delineate biphasic insulin secretion, we separated the first-phase of insulin release by fasting mice for 16 hours and injecting them with glucose, then collecting blood and reading glucose levels at short time intervals: 1, 5, 15, 30, and 60 minutes. NPKO mice had a defect in the first-phase insulin release, with significantly less insulin produced 5 minutes after glucose injection than controls (**Figure 4F**).

Moreover, analysis of single-cell patch-seq data shows that expression of NFIA is significantly correlated to exocytosis (normalized to Ca^2+^ influx) and sodium conductance in beta cells from non-diabetic donors (**Figure 4G**) [https://doi.org/10.1101/555110].

Patch-seq data also confirms that *NFIA* is heterogeneously expressed in beta cells, as we observed from our immunohistochemical staining (**Figure 1**). These findings demonstrate that NPKO pancreatic beta cells are dysfunctional and indicate that there is a defect in the recruitment, docking, or exocytosis of the ready releasable pool of insulin granules.

*NFIA directly regulates Rab39b to modulate granule recruitment and insulin secretion* To better understand how NFIA, a transcription factor, could regulate insulin secretion, we isolated islets from control and NPKO mice and performed RNA-sequencing (**Figure 5A**). RNA-sequencing revealed significant changes in 426 genes (*q*<0.05) after deletion of NFIA, with 291 decreased and 135 increased (**Figure 5B**). Gene ontology analysis of significantly decreased genes showed processes associated with pancreatic secretion, transport, cell adhesion, insulin secretion, and digestion, among others (**Figure 5C**).

**Figure 5.**
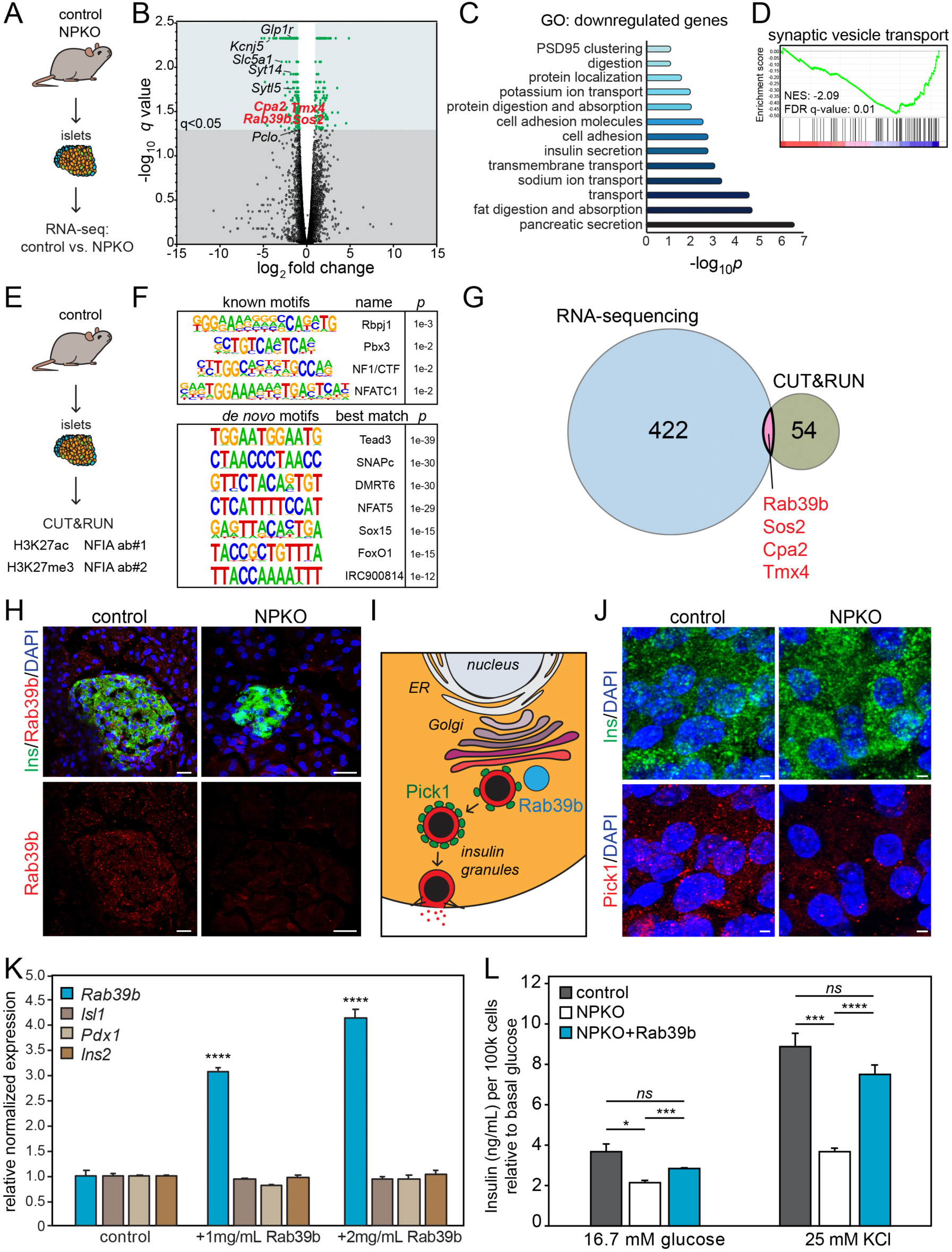
NFIA directly targets Rab39b to regulate insulin granule trafficking in pancreatic beta cells. A. Schematic of RNA-sequencing from control and NPKO mouse islets. B. Volcano plot showing transcriptional changes uncovered by RNA-sequencing of control and NPKO adult mouse islets. The change in expression is shown on the x-axis while the significance is plotted on the y-axis. Select transcripts are shown, with the four confirmed by CUT&RUN analysis in red. N=2 islet preps from 4-6 mice per genotype. C. Gene ontology (GO) analysis of transcripts significantly decreased (*q<*0.05) in NPKO islets. D. Gene set enrichment analysis shows synaptic vesicle transport significantly decreased in NPKO islets compared to control islets. N=3 mice per genotype. Normalized enrichment score (NES) = -2.09, FDR q-value = 0.01. E. Schematic of CUT&RUN from control and NPKO mouse islets. Two NFIA antibodies and one antibody for H3K27ac, H3K27me3, and Nkx6.1 were used. N=4 mice per genotype. F. Motif enrichment of NFIA binding sites showing known and *de novo* motifs. G. Venn diagram comparing transcripts significantly changed in NPKO islets from RNA-sequencing to regions directly bound by NFIA using both antibodies from CUT&RUN analysis. Four genes were shared between the datasets. H. Immunofluorescent staining of adult control and NPKO pancreas showing beta cells (Ins in green) with Rab39b in red. Nuclei are marked by DAPI in blue. Scale bars= 20 μm. I. Schematic showing Rab39b association with Pick1, and Pick1 association with insulin granules. J. Immunofluorescent staining of adult control and NPKO beta cells, with Ins shown in top in green and Pick1 shown in the bottom in red. Scale bars= 2 μm. K. Quantitative PCR of ectopic *Rab39b* expression at two different concentrations, 1 mg/mL and 2 mg/mL, in mouse islets, relative to *Gapdh* and normalized to control. Error bars show SEM, *\*\*\*\*p*<0.001. L. Phenotypic rescue of NPKO islets. NPKO islets were transfected to express Rab39b for 6-days before glucose stimulated insulin secretion assays were performed on control, NPKO, and NPKO+Rab39b islets. Insulin secretion was determined by ELISA and normalized to cell number relative to insulin secretion after 2.8 mM glucose stimulation. N=4 mice per genotype. Error bars show SEM, \*\*\*\**p<*0.001, \*\*\**p<*0.005, \**p<*0.05, *ns*=not significant between mice.

Gene set enrichment analysis (GSEA) showed significant changes in processes including synaptic vesicle transport, vesicle priming, and digestion (**Figure 5D, S5A**). To find direct binding sites of NFIA in islets, we used CUT&RUN target analysis (Skene and Henikoff, 2017) (**Figure 5E**). Using two different NFIA antibodies in biological quadruplicate, we found 53 genes with peaks consistent between all eight conditions and enrichment for NF1/CTF motifs in bound regions, as expected, among other sites that may indicate co-factor binding sites such as Rbpj1 and Tead3 (**Figure 5F**). In addition, we profiled marks of active (H3K27ac) and inactive (H3K27me3) chromatin in primary mouse islets, which showed both marks centered around NFIA binding sites, supporting a role for NFIA in regulating gene expression in adult endocrine cells (**Figure 5C**), as we also observed by RNA-sequencing (**Figure 5B**).

To uncover the top transcriptional targets of NFIA, we cross-referenced the 426 differentially expressed transcripts identified by RNA-sequencing with the 53 putative NFIA binding sites by CUT&RUN, reducing the list to four genes: Tmx4, Sos2, Rab39b, and Cpa2 (**Figure 5G**). After loss of NFIA, *Rab39b* expression was significantly decreased. Rab39b, a Rab GTPase, forms a complex with Pick1 on secretory vesicles in hippocampal neurons to regulate alpha-amino-3-hydroxy-5-methyl-4-isoxazole propionic acid receptor (AMPAR) trafficking to the membrane (Mignogna et al., 2015). This process occurs through Pick1 binding to islet cell autoantigen 69-kD (ICA69) to regulate early exocytosis before switching to Pick1 only binding in more mature vesicles (Cao et al., 2013). We immunostained the adult control and NPKO pancreas and found decreased Rab39b protein expression in beta cells (shown by Ins, **Figure 5H, S5F**).

Similarly to the role of Rab39b in neurons, RNA-sequencing showed significantly decreased expression of the AMPAR gene *Gria2*. Immunostaining for AMPAR (Gria2) confirmed decreased expression, however, we found Gria2 was not expressed in beta cells but was restricted to insulin-negative alpha cells on the periphery of endocrine islets (**Figure S5D**). In addition, we observed no change in islet structure as illustrated by immunostaining for phalloidin (marking F-actin), Tubulin, E-cadherin (Cdh1) or for synaptic vesicle docking protein Syntaxin-3 (Stx3) (**Figure S5E, S5F**). Collectively we demonstrate that NFIA directly regulates expression of Rab39b, which is known to regulate vesicular trafficking in multiple contexts (Gambarte Tudela et al., 2019; Mignogna et al., 2015; Shi et al., 2017).

Loss of Pick1 in pancreatic beta cells causes similar defects in blood glucose homeostasis and insulin secretion as we found after loss of NFIA; additionally, the recruitment and docking of insulin granules follows a similar mechanism to AMPAR trafficking (Cao et al., 2013). However, how Rab39b is associated with this process in the pancreas is currently unknown. From our data, we propose a model in which NFIA directly regulates *Rab39b* expression, with Rab39b then interacting with Pick1 to promote Pick1 association with insulin granules for exocytosis (**Figure 5I**). Indeed, in NPKO beta cells, which have a decreased level of Rab39b, we found altered localization of Pick1 with few isolated punctae, whereas punctae are spread among insulin granules in control beta cell (**Figure 5J, S5G**).

Finally, to determine whether the NFIA-Rab39b axis was responsible for the defects in insulin secretion in NPKO beta cells, we re-expressed Rab39b by transfection of a plasmid driving expression of Rab39b in control and NPKO mouse islets, and confirmed a 4-fold increase by qPCR (**Figure 5K**). We then performed GSIS in NFIA-deficient islets and found that expressing Rab39b in NPKO islets restored glucose responsiveness and secretion after depolarization with KCl, rescuing the phenotype from loss of NFIA (**Figure 5L**). Together, these studies show that NFIA regulates *Rab39b* expression, which is required for Pick1 association with insulin granules and proper insulin granule recruitment and docking to the membrane. These results shed light on the mechanisms regulating insulin secretion in pancreatic beta cells, and uncover a transcription factor that governs this process.

## Discussion

In this study, we found that NFIA regulates pancreatic physiology through modulation of insulin and zymogen granule trafficking. In the absence of NFIA, mice developed elevated blood glucose levels and defects in GSIS due to impairments in beta cell function. In addition, impaired zymogen granule trafficking led to an accrual of granules within acinar cells and an inability to break down lipids. In NPKO mice we observed accumulation of insulin granules within pancreatic beta cells and fewer granules docked in the readily releasable pool. This accumulation led to a higher level of insulin in the pancreas of NPKO compared to control mice but less insulin circulating in the blood stream. Furthermore, isolated NPKO islets did not respond to glucose, KCl stimuli, or numerous alternative insulin secretagogues.

We previously identified NFIA in the developing pancreas, and found that NFIA regulates trafficking related genes at embryonic day 17.5, including Notch-related trafficking gene Mib1 (Scavuzzo et al., 2018). In this study, we describe how NFIA transcriptionally regulates trafficking genes including Rab39b in the adult pancreas, thus impacting the physiology of secretory cells. Rab39b is a small GTPase involved in vesicle trafficking, and its loss of function in neurons leads to a number of defects including Parkinson’s disease, autism, epilepsy, and macrocephaly (Giannandrea et al., 2010; Shi et al., 2017; Wilson et al., 2014). NFIA-related disorders lead to similar defects, including epilepsy, macrocephaly, developmental delays, and cognitive impairments (Ji et al., 2014; Koehler et al., 2010; Labonne et al., 2016; Mandal et al., 2015). The mechanism of Rab39b in the pancreas has not been established, however in neurons Rab39b forms a complex with Pick1 to facilitate the trafficking of AMPA receptor subunit GluA2 to the membrane (Mignogna et al., 2015). In pancreatic beta cells, Pick1 deletion leads to hyperglycemia and diabetes as we observed after loss of NFIA (Cao et al., 2013; Herlo et al., 2018; Li et al., 2018). This study supports that insulin granule secretion follows a similar mechanism to neurons, and is the first to establish a role of NFIA as well as Rab39B in the adult pancreas.

As NFIA is expressed in other pancreatic secretory cells including acinar cells, we also examined the effect of NFIA on acinar physiology. We observed increased amylase activity in the pancreas of NPKO mice but not in blood serum, suggesting that trafficking defects affected other secretory cells that express NFIA. NPKO mice exhibited accumulation of zymogen granules, decreased circulating lipids, and retention of fat in the stool of animals, showing impaired digestion. The defects in secretion could lead to precocious activation of zymogens in the pancreas, leading to necrosis and pancreatitis, and we indeed observed fibrosis in the NPKO pancreas and a massive increase in amylase activity in blood serum after caerulein injection. Loss of NFIA thus impairs trafficking of granules not only in endocrine cells but also in secretory acinar cells in the pancreas, causing digestive deficiencies and pancreatic fibrosis. Interestingly, we observed increased granule size and defects in amylase secretion; previously, different Rab GTPases have been implicated in regulating zymogen granule size and amylase secretion (Hou et al., 2016). Whether NFIA regulates acinar secretion through the same mechanism as beta cells will be interesting to understand. It will also be intriguing to harness inducible Cre lines such as the acinar specific Ptf1a-cre^ERt2^ to further investigate the role of NFIA in acinar cells, and to isolate acinar cells to determine whether the mechanism of dysfunction is distinct from NFIA-Rab39b in endocrine beta cells.

NFIA is known to regulate cell fate in the central nervous system and myeloid cells (Glasgow et al., 2014; Hiraike et al., 2017; Kang et al., 2012; Mei et al., 2016; Tian et al., 2015; Yang et al., 2018), however, the adult functions of NFIA in these cells have not yet been established. Here, we illustrate a new function of NFIA not related to cell cycle, cell fate determination, or organ formation. Instead, we found that NFIA regulates trafficking in the adult pancreas. Interrogating other cell types in which NFIA is expressed may shed light on whether it similarly plays a role in regulating trafficking, secretion, and endocytosis. Loss of NFIA in the pancreas leads to inadequate insulin release, resulting in prolonged hyperglycemia. Understanding the mechanism of NFIA in regulating exocytosis will provide insight into the functioning of pancreatic beta cells, linking a transcription factor to the regulation of hormone secretion. Further, as type 2 diabetes continues to rise in prevalence, delineating the pathways affecting insulin exocytosis and contributing to hyperglycemia is important to discovering new treatment strategies for the disease.

## Acknowledgements

We thank Katrina Wamble, Abraham Akerele, and Robert Sharp for superb technical work. Conditional NFIA mice and NFIA homemade antibodies were generously provided by Dr. Benjamin Deneen (Baylor College of Medicine). We thank Dr. Hugo Bellen for his generosity to share his transmission electron microscope for this work. We thank Dr. Paul Tesar for his time and support. We thank Kevin Allan, Erin Cohn, Mayur Madhavan, and Andrew Scavuzzo for valuable feedback and helpful discussions. This work was supported by the NIH (P30-DK079638 to M.B., 5T32HL092332-13 to M.A.S. and M.B.), the McNair Medical Foundation (to M.B.), the New York Stem Cell Foundation (to M.A.S.), the confocal core at the BCM Intellectual and Developmental Disabilities Research Center (NIH U54 HD083092 from the Eunice Kennedy Shriver National Institute of Child Health and Human Development). J.C.K was supported by T32GM008231. A.C. was supported by R01DK115454, by the Ted Nash Long Life Foundation, and is a CPRIT Scholar in Cancer Research (RR140038).

## Author Contributions

Conceptualization, M.A.S., J.C., and M.B. Methodology, M.A.S., J.C., D.Y., M.C.H., A.R.W., J.C.S., X.D., L.D., and J.C.K. Investigation, M.A.S., J.C., D.Y., M.C.H., A.R.W., J.C.S., X.D., L.D., and J.C.K. Resources, A.C., S.R.Q., P.E.M., and M.B. Writing –Original Draft, M.A.S. and M.B. Writing – Review & Editing, M.A.S. and M.B.

## Supplemental Figures

**Figure S1. NFIA is expressed in the adult pancreas.**

A. LocusZoom plot of the p-values for variants in *NFIA* in the GERA meta-analyses for high-density lipoprotein cholesterol levels (Hoffman et al. 2018). Variants displayed fall within the gene boundaries of *NFIA* or within 500 kilobases on these gene boundaries. P-values were previously generated in meta-analyses of the GERA cohort (Hoffman et al. 2018). Each ancestry group in the GERA cohort were analyzed individually and meta-analyzed together. The lead SNP (rs55878063, p = 1.5 × 10^−8^) is highlighted in purple. LD estimates are not displayed because the discovery cohort was multi-ethnic. Summary statistics were downloaded from the GWAS Catalog (Study: GCST007140).

B. Western blot of NFIA expression in the whole pancreas and in isolated islets. B-actin is shown as a loading control.

C. Quantitative PCR of human *NFIA* expression in the pancreas at fetal week 20 and in adult, relative to *GAPDH*.

D. Immunofluorescent staining of d20 control and NPKO pancreas showing beta cells (Ins in green) with ducts (DBA in red). Nuclei are marked by DAPI in blue. Scale bars=100 μm.

E. Body weight of 3 month old (3M) mice, separated by sex; N=3 mice per genotype.

F. Dissected pancreas from adult control and NPKO mouse.

G. Pancreas to body weight ratio of control and NPKO mice. *p*<0.001, N=11 NPKO and N=12 control mice.

**Figure S2. Loss of NFIA leads to defects in blood glucose homeostasis and lipid breakdown.**

A. Random blood glucose levels in control and NPKO mice. N=5 control and N=3 NPKO mice. \**p<*0.05.

B. Random blood glucose levels in control and NFIA^fl/fl^; Ngn3-cre mice. N=30 control and N=29 NPKO mice. \*\*\*\**p<*0.001.

C. Random blood glucose levels in control and NFIA^fl/fl^; Ins2-cre mice. N=12 mice per genotype. \*\*\**p<*0.005.

D. Insulin tolerance tests were performed by injecting mice fasted for short duration (6 hours) with 0.75 IU insulin/g body weight and measuring blood glucose levels. N=4 control and N=3 NPKO mice. *\*\*\* p*< 0.005.

E. Area under the curve for insulin tolerance test showing no change.

F. Fresh stool from control and NPKO mice was collected and directly put into water.

**Figure S3. Accumulation of granules within pancreatic cells.**

A. Increased zymogen granule size in area (μm^2^) in adult NPKO mouse pancreatic acinar cells as seen in Figure 4C. N=4 control and N=2 NPKO mice, n=893 control granules and n=835 granules in NPKO. \*\*\*\**p<*0.001.

B. Amylase activity in control and NPKO circulating blood serum as U/L. N=2 control and N=3 NPKO mice.

C. Caerulein was injected via IP at 50 µg/kg body weight of the individual mouse. A total of six injections were given, one dose per hour for each mouse, and tissue collected at the end of treatment for H&E staining. Each image is from an individual mouse. Scale bars = 100 μm.

D. Caerulein was injected via IP at 50 µg/kg body weight of the individual mouse. A total of six injections were given, one dose per hour for each mouse, before amylase activity was measured in blood serum. N=6 control and N=5 NPKO mice, error bars show SEM. \**p*<0.05, \*\*\**p<*0.005, \*\*\*\**p*<0.001 compared to controls treated with saline. #*p*<0.05, ###*p*<0.005, ####*p*<0.001 compared to controls treated with caerulein.

E. Percentage of mature insulin vesicles as determined by dense core vesicle versus light vesicles.

**Figure S4. Deletion of NFIA does not impair cellular respiration.**

A. Schematic diagram of the immediate, first phase insulin secretion and the slower, second phase insulin secretion.

B. Glucose stimulated insulin secretion assay normalized to control 2.8 mM glucose. N=8 mice per genotype, n=2 technical replicates. \**p<*0.05.

C. Baseline insulin secretion in KREBS media unchanged between control and NPKO mice (see Figure 3E). N=4 mice per genotype.

D. Transmission electron microscopy of mitochondria in control and NPKO mouse pancreatic cells. Scale bars = 400 nm.

E. Mitochondria size in area. N=2 mice per genotype, n=81 control and n=61 NPKO mitochondria assessed.

F. Distribution of mitochondria area binned into 100 nm size ranges.

G. Number of mitochondria normalized to cytoplasmic area relative to control. N=3 control and N=2 NPKO mice; n=14 control and n=5 NPKO cells.

H. Oxygen consumption rate (OCR) of dissociated control and NPKO islet cells. N=3 mice per genotype, n=3 technical replicates.

I. Extracellular acidification rate (ECAR) of dissociated control and NPKO islet cells. N=3 mice per genotype, n=3 technical replicates.

**Figure S5. NFIA regulates granule exocytosis.**

A. Gene set enrichment analysis shows synaptic vesicle priming significantly decreased in NPKO islets compared to control islets. N=3 mice per genotype.

B. Gene set enrichment analysis shows digestion decreased in NPKO islets compared to control islets. N=3 mice per genotype.

C. H3K27ac and H3K27me3 histone marks centered around NFIA peaks. ChIP-seq fragment depth shows enrichment of both repressive and active marks at NFIA binding sites. N=4 control mice per ChIP-seq experiment.

D. Immunostaining of the AMPAR, Gria2 (in red), surrounding beta cells (Ins, in green). Cdh1 marks the epithelium in white while DAPI marks nuclei in blue. Scale bars = 50 μm.

E. Immunostaining of the F-actin by phalloidin (in green), Tubulin (in red), and the epithelium by Cdh1 (in white). DAPI marks nuclei in blue. Scale bars = 50 μm.

F. Immunostaining of the synaptic vesicle docking protein Stx3 (in red) proximal to beta cells (Ins, in green). The Rab GTPase Rab39b is shown in white while DAPI marks nuclei in blue. Scale bars = 2 μm.

G. Immunostaining of Pick1 (in red) co-localized in control beta cells with insulin granules (Ins, in green). DAPI marks nuclei in blue. Scale bars = 2 μm.

## Experimental Procedures

### Animals

Animal studies were approved by the Baylor College of Medicine Institutional Animal Care and Use Committee. Mice were housed at 22–24°C with a 12 hour light/12 hour dark cycle with standard chow (Lab Diet Pico Lab 5V5R, 14.7% calories from fat, 63.3% calories from carbohydrate, 22.0% calories from protein) and water provided ad libitum unless otherwise indicated. All mice studied were on a mixed background (129SvEv, FVB, and C57BL/6J).

NFIA conditional knockout mice were generated as described previously (Scavuzzo et al. 2018). Both female and male animals were used in this study. The Pdx1-cre mice were obtained from The Jackson Laboratory (JAX 024968) and originally made by Gu et al. (2002). The Ngn3-cre mice were obtained from The Jackson Laboratory (JAX 06333), and originally made by Schonhoff et al. (2004). The Ins2-cre (RIP-cre) mice were obtained from The Jackson Laboratory (JAX 03573) and originally made by Postic et al. (1999).

All genotyping was done using the HotStart Mouse Genotyping Kit with its instructed PCR reaction set up from KAPA Biosystems. For the NF1A^fl/fl^ genotype, two primers were used (F-CCGAGCATGGAGATTTGCTTTG R-TGTGCGGCCTACTTCCCAGA) with an annealing temperature of 55°C, resulting in a wildtype band of 511 bp and a floxed band of 544 bp. To genotype Cre, two primers were used (F-GAACGCACTGATTTCGACCA R-AACCAGCGTTTTCGTTCTGC) with an annealing temperature of 53°C, resulting in one band of 325 bp.

### Publicly available genome-wide association studies (GWAS)

Summary statistics from the phs000221.v1.p1 study in dbGAP (NCBI dbGAP analysis accession: pha003059) were downloaded and visualized with LocusZoom (locuszoom.org) with an emphasis on a suggestive variant for diabetes (rs11585396, p = 6.18 x 10^-6^) upstream of *NFIA*. These p-values were previously generated in a genome-wide association study of individuals taking medication for diabetes or glucose in the National Heart, Lung and Blood Institute (NHBLI) Family Heart Study (FamHS) at Visit 1 (NCBI dbGAP analysis accession: pha003059).

The NHBRI-EBI GWAS Catalog (Buniello, et al. 2019) was queried for significant (p < 5 x 10^-8^) or suggestive (p < 1 x 10^-5^) evidence of genetic associations in *NFIA* with HDL levels. Summary statistics were downloaded from the GWAS Catalog for the GCST007140 study (Hoffman et al. 2018). P-values for variants in *NFIA* or within 500 kilobases of the gene boundaries were visualized in LocusZoom (locuszoom.org). This study included summary statistics from meta-analyses of the multi-ethnic Genetic Epidemiology Resource on Adult Health and Aging (GERA) cohort (Hoffman et al. 2018) in which each ancestry group was analyzed individually and in combinations. This included 76,627 Non-Hispanic white, 7,795 Latino, 6,855 East Asian, 2,958 African American, and 439 South Asian individuals (Hoffman et al. 2018). In their study, they identified a genome-wide significant variant for HDL cholesterol levels in *NFIA* (rs55878063, p = 1.5 x 10^-8^). This variant also exhibited a suggestive association with triglyceride levels (p = 1 x 10^-6^) in the GCST007142 study (Hoffman et al. 2018) in the GWAS Catalog.

Summary statistics variants within *NFIA* +/- 500 kilobases were downloaded from the GeneATLAS for diabetes (UK Biobank field: 20002; Field codes: 1220, 1223, 1521, 1221, 1222) and visualized with LocusZoom (locuszoom.org). GeneATLAS is a publicly available repository of genetic associations in the UK Biobank (Canela-Xandri et al. 2018). The GWAS of diabetes in GeneATLAS was performed on 21,105 diabetes cases and 431,159 controls. Although no genome-wide significant variants were identified in *NFIA* for this trait, there was a suggestive association for diabetes in *NFIA* (rs1984296, p = 3.77 x 10^-6^).

### Immunostaining and Western blot analysis

Whole pancreas was fixed in 4% paraformaldehyde at 4°C overnight, washed and embedded in O.C.T. compound (TissueTek, Sakura Finetek) for immunofluorescence or paraffin for immunohistochemistry and performed as described (Fancy et al., 2012).

Briefly, after deparaffinization, tissue was subjected to antigen retrieval for 7 minutes in 1x sodium citrate pH6.0, followed by 15 minutes of 3% H_2_O_2_ incubation, blocking and finally incubated with NFIA antibody. Following washes, slides were incubated with ImmPRESS^TM^ HRP anti-rabbit (Vector Laboratories). Colorimetric reaction was done using ImmPACT DAB peroxidase (Vector Laboratories). Slides were then counterstained with Harris Hematoxylin (Poly Scientific R&D), re-hydrated and mounted using Permount (Fisher chemicals).

Protein lysates from whole pancreas were prepared in 1 mL of RIPA buffer supplemented with phosSTOP phosphatase and protease inhibitors (Roche). Protein concentrations were measured using the BCA kit (Thermo Scientific) and then diluted in 4x Laemmeli buffer (40% glycerol, 8% SDS, 240 mM Tris-HCl pH 6.8, 5% b-mercaptoethanol, 12.5mM EDTA, 0.04% bromophenol blue) and RIPA buffer at 10 μg/uL. 10 μL of lysate was resolved on 8% SDS-PAGE and transferred to Nitrocellulose membrane (BioRad). Membranes were blocked with 5% milk in Tris-buffered saline with 1% Tween 20 (TBST) for 1 hr, followed by overnight incubation at 4°C with primary antibodies. Membranes were washed three times with TBST before incubation for 1 hr with secondary antibodies. Expression was quantified using ImageJ and normalized to β-actin.

For immunofluorescent staining, slides were blocked for 30 minutes room temperature with PBS+0.1% Triton X-100 (VWR) with 5% donkey serum (Jackson ImmunoResearch) then incubated overnight at 4°C in primary antibodies. Slides were washed, incubated with secondary antibody for 1 hour at room temperature, and washed. All secondary antibodies were purchased from Jackson ImmunoResearch. Slides were mounted in Fluoromount G (SouthernBiotech), covered with coverslips, and sealed with nail polish. Nuclei were stained with DAPI (Invitrogen). Imaging was performed on Leica CTR DM6000 FS, Leica TCS SPE, and Zeiss Axioplan 2.

### Histology

Whole pancreas was fixed in 4% paraformaldehyde 2-16 hours, washed, incubated in 30% sucrose overnight at 4C, and embedded in O.C.T. compound for immunofluorescent staining or dehydrated in ethanol series and embedded in paraffin for H&E and Masson’s Trichrome staining. H&E staining were performed as described (Towers, 1953; Wu, 1940), and Masson’s Trichrome staining was performed using Masson’s Trichrome reagents from Sigma (HT15).

### Metabolic analysis

For in vivo assays, blood samples were collected by tail vein nick or by cutting <0.5 mm of the tail tip for serum insulin (Mouse Insulin ELISA, ALPCO Diagnostics) and glucose (glucometer, Contour next, Bayer) measurements, respectively. Control and NFIA^fl/fl^;Pdx1-cre adult (7-10 weeks) animals blood glucose levels were measured randomly, during a short duration fast (7 hours), or long duration fast (16 hours). Glucose tolerance tests (GTTs) were performed in mice fasted for 16 hours injected intraperitoneally with 2 mg D-glucose/g body weight. Insulin tolerance tests were performed by injecting mice fasted for short duration (6 hours) with 0.75 IU insulin/g body weight and measuring blood glucose levels. Pancreatic insulin content measurements were taken by acid-ethanol extraction to extract protein from pancreas. Insulin content was measured using Mouse Insulin ELISA kit (ALPCO diagnostics). Amylase activity levels in the blood serum and pancreas were determined using Amylase activity assay (Sigma). Blood plasma was collected from adult NPKO and control mice for lipid panel analysis. Different lipoprotein particles in plasma were analyzed at the Mouse Metabolic Core at Baylor College of Medicine using the FPLC system from Amersham Pharmacia Biotech. In the FPLC system, 2 Superose 6 columns used for plasma fractionation are connected in series. Using Unicorn software, complete separation process is controlled by computer. It separates the lipoproteins into very low density (VLDL), low and intermediate density (IDL & LDL) and high-density lipoproteins (HDL). A total of 250 μl of plasma sample is separated using an elution buffer containing 1 mM EDTA, 154 mM NaCl, and 0.05% NaN3, pH 8.2 in 44 fractions of 0.5 ml each. Cholesterol and triglyceride concentrations in individual fractions are determined with commercial kits and plotted to compare the peak height and shift.

### Caerulein challenge to acinar cells

Caerulein was injected into control and NPKO mice via IP at 50µg/kg body weight of the individual mouse injected at a 10 μg/mL volume. A total of six injections were given, one dose per hour for each mouse. At the end of the six hours, blood was collected to extract serum at different timepoints. Amylase activity levels in the blood serum and pancreas were determined using Amylase activity assay (Sigma). Serum samples were not diluted, and 5 μL was used per well. For tissue, 50mg of tissue was homogenized in 0.5 mL of amylase buffer before diluting 1 to 100 and using 1 μL per well. All samples were run in technical duplicates.

### Islet preparation

Mouse islet isolation was performed by common bile duct perfusion with ice cold 0.8 mM Collagenase P (Roche), dissection, additional gentle dissociation with Collagenase P and further purification in Histopaque gradient (Sigma) as described previously (Szot et al., 2007).

### Glucose dynamics ex vivo in islets

Mouse islets were washed with Krebs buffer (126 mM NaCl, 2.5 mM KCl, 25 mM NaHCO_3_, 1.2 mM NaH_2_PO_4_, 1.2 mM MgCl_2_, and 2.5 mM CaCl_2_) and preincubated in Krebs buffer with 0.1% BSA for 2 hours at 37°C to remove residual insulin. Media was collected after preincubation and after each subsequent incubation. Islets were incubated in low-glucose (2.8 mM), high-glucose (16.7 mM), or 25 mM KCl in KREBS with 0.1% BSA to induce depolarization and insulin dumping. Islets were then collected and dispersed into single cells using TrypLE Express to measure cell number for normalization using Countess Automated Cell Counter (Invitrogen). Insulin content was determined using Mouse Insulin ELISA kits (ALPCO Diagnostics).

### Response to secretagogues

Mouse islets were washed with Krebs buffer and preincubated in Krebs buffer with 0.1% BSA for 2 hours at 37°C to remove residual insulin. Media was collected after preincubation and after each subsequent one-hour incubation. Islets were incubated in 20 nM Exendin-4 (Enzo Life Sciences ENZ-PRT11-0500), 100 μM IBMX (Enzo Life Sciences BLM-PD140-0200), 1 μM Forskolin (Enzo Life Sciences BML-CN100-0010), 20 μM PMA (company), or 100 mM Diazoxide (Sigma Aldrich D9035) with 2.8 mM glucose or 30 mM KCl in KREBS with 0.1% BSA. Islets were then collected to measure DNA content for normalization. Insulin content was determined using Mouse Insulin ELISA kits (ALPCO Diagnostics).

### Patch-seq analysis

Analysis of patch-seq data from human beta-cells is performed using single-cell count tables and electrophysiological annotations from [https://doi.org/10.1101/555110]. Total sequencing counts are normalized to counts per million (cpm) and transformed to log2 values after addition of a pseudocount. Outliers in electrophysiology are removed by quantile filtering of the highest and lowest 3% of cells for each parameter, followed by log transformation and scaling (0-1 range). We then used the set of 484 genes with high correlation to electrophysiological parameters reported in [https://doi.org/10.1101/555110] to perform a tSNE projection of all beta-cells from non-diabetic donors (glucose: 5-10mM). Scaled gene expression values (for NFIA) and normalized values of electrophysiological parameters that show significant correlation to NFIA expression (i.e. exocytosis normalized to Ca^2+^ and Na^+^ conductance) are shown.

### Seahorse assay

We assessed the respiratory capacity of NPKO and control cells using the Cell Mito Stress Test Kit (Agilent Cat. #103010-100) on the Agilent Seahorse XFp instrument. The day prior to the experiment, one XFp Extracellular Flux Cartridge (Agilent Cat. #102983) was hydrated with Agilent Seahorse XF Calibrant (Agilent Cat. #103059) and stored overnight in a humidified 37°C non-CO2 incubator. One Agilent cell culture miniplate (Agilent Cat. #102984) was coated with Corning Cell-Tak Cell and Tissue Adhesive (Corning Cat. #354240) and stored at 4°C following the vendor’s protocol. On the day of the assay, NPKO and control islets were dissociated, and the cells were washed and resuspended in Agilent Seahorse XF base medium (Agilent Cat. #103193) supplemented with 1mM pyruvate (Thermo Fisher Cat. #11360070), 2 mM L-glutamine (Thermo Fisher Cat. #25030), and 10 mM glucose (Sigma Aldrich Cat. #G8769). Cells were quantified with a Nexcelom Cellometer Auto 2000 and diluted to a final concentration of 1×10^6 cells/mL in supplemented Agilent Seahorse medium. Three wells of the coated cell culture mini plate were then loaded with 50 μL each of the control cells, and three wells with 50 μL each of the NPKO cells, for a final concentration of 50,000 cells/well. The first and final wells were loaded with 50 μL of medium. The plate was centrifuged for one minute at 200 x g with no braking to facilitate cell adhesion. The plate was then placed in a 37°C non-CO2 incubator for 25 minutes. 125 μL of Seahorse medium was added to each well, and the plate was incubated for an additional 25 minutes. During these incubations, the oligomycin, FCCP, and rotenone/antimycin A stocks were rehydrated in supplemented Seahorse medium and diluted to form working solutions. 25 μL of oligomycin was loaded into Port A of each sensor on the hydrated cartridge, 25 μL of FCCP was loaded into Port B of each sensor, and 25 μL of rotenone/antimycin A was loaded into Port C of each sensor. After loading the cartridge and calibrating the instrument, the cell plate was loaded, and the Mito Stress Test assay was run using final concentrations of 1.0 μM oligomycin, 0.5 μM FCCP, and 0.5 μM rotenone/antimycin A.

### Quantitative PCR

Total RNA was isolated by phenol-chloroform extraction using TRIzol (Thermo Fischer Scientific). Genomic DNA was depleted by DNAse treatment (Invitrogen) before reverse transcription using iScript cDNA Synthesis kit (BioRad). Real-time PCR was performed with diluted cDNAs in a 15 μL reaction volume using Kapa SYBR Fast qPCR mix in duplicates and analyzed using the CFX96 Touch Real-Time PCR Detection System and computing ΔΔCq using the CFX96 software. Genes were normalized to *Beta-actin* and *Gapdh*.

### Transmission electron microscopy

Modified Karnovsky’s fixative was buffered to 320 mOsmol/kg with sodium Cacodylate buffer. Adult mice were collected, weighed and given isoflurane as a sedative. After surgical opening, the vascular system was flushed with normal saline before perfusion with the buffered fixative. The pancreas was dissected and sliced into 1 mm slices.

Tissue was placed into a scintillator vial with buffered fixative and placed on a rotator overnight. Three days later the tissue was processed inside a Ted Pella Bio Wave Vacuum Microwave. Samples were fixed again, followed by 3x sodium cacodylate buffer rinses, post-fixed with 1% buffered osmium tetroxide, and followed again with 3 millipore water rinses. Ethanol concentrations from 30-100% were used as the initial dehydration series, followed with propylene oxide as a final dehydrant. Samples were gradually infiltrated with 3 propylene oxide and Embed 812 resin graded ratios into 3 changes of pure resin under vacuum. Samples were allowed to infiltrate in pure resin overnight on a rotator. The samples were embedded into regular Beem capsules and cured in the oven at 62°C for 5 days. The polymerized samples were sectioned at 50 nm. Grids were then stained with 1% uranyl acetate for fifteen minutes followed by lead citrate for three minutes before TEM examination. TEM images were captured using a JEOL JEM 1010 transmission electron microscope with an AMT XR-16 mid-mount 16 mega-pixel digital camera.

### Morphometric analysis

Docked insulin granules were measured by counting the number of granules touching the plasma membrane normalized to the total number of granules per beta cell, in 4 beta cells per group. Distance from the plasma membrane to the center of insulin granules was measured and binned into 5 groups (0-100 nm, 100-250 nm, 250-500 nm, 500-1000 nm, and greater than 1000 nm). Data is shown as percentage of granule density (granules/binned area) relative to cytoplasmic area (area of beta cell cytoplasm). The total number of insulin granules per cytoplasmic area was also measured. Zymogen granule number was calculated by counting the number of zymogen granules per acinar cluster, counting a total of 14 control and 10 NPKO clusters. Zymogen granule size was counted by calculating the area of each granule in ImageJ. 893 control and 835 NPKO granules were quantified. The size of zymogen granules was binned into 6 groups (0-200 nm, 200-400 nm, 400-600 nm, 600-800 nm, 800-1000 nm, and greater than 1000 nm). The distribution of zymogen granules within an acinar cluster was measured as the area of granules in the cytoplasm normalized to the total area of the cluster. Distribution maps were created by tracing over clusters and granules in Illustrator. Mitochondria number was counted and normalized to cytoplasmic area, and mitochondria size was determined in area (nm^2^) for 3 control and 2 NPKO mice. Dense and light core insulin granules were counted to determine the percentage of mature insulin granules in 3 control and 2 NPKO mice.

### RNA-sequencing

RNA was isolated from adult islets from control and NPKO by phenol-chloroform extraction using TRIzol. Quality RNA was determined by analysis by formaldehyde gel and Bioanalyzer (Agilent) before cDNA synthesis using the TruSeq Stranded mRNA kit. Paired-end RNA-sequencing was performed using the Illumina NextSeq500 platform for 150 cycles as a depth of 130M reads (NextSeq500 Mid Output Kit). Sequencing reads were aligned to the mouse genome (RefSeq mm10) using TopHat Alignment (version 1.0.0). Gene expression was quantified by FPKM. Differential expression between groups of samples was computed by Cufflinks Assembly & DE (version 1.1.0). NFIA-dependent genes were identified as genes differentially expressed (q<0.05) between the control and NPKO pancreas (fold change > 1.5 and < 0.75) with p<0.05 for validated expression changes. GSEA 3.0 was used for gene set enrichment, and The Database for Annotation, Visualization and Integrated Discovery v6.7 (Dennis et al., 2003) was used for GO analysis and KEGG pathway analysis for NFIA-dependent genes.

### CUT&RUN

CUT&RUN experiments were carried out as described (Skene et al., 2018). Briefly, 200,000 islet cells were washed in wash buffer (20 mM HEPES, pH7.5, 150 mM NaCl, 0.5 mM Spermidine and complete protease inhibitor (EDTA-free, Roche), captured with Concanavalin A beads (Polysciences, Warrington, PA) and incubated with primary antibodies overnight at 4°C. After washing with Dig-wash buffer (20 mM HEPES, pH7.5, 150 mM NaCl, 0.5 mM Spermidine, 0.08% Digitonin and protease inhibitors), cells were resuspended in 50 μL Digwash buffer and 2.5 μL of protein A-MNase (1:10 diluted, batch 6 from Steve Henikoff) and incubated at room temperature for 10 minutes. Cell pellets were washed again and placed in a 0°C metal block, and 2 mM of CaCl2 was added and incubated for 45 minutes. MNase reaction was terminated by the addition of 2XSTOP buffer and incubated at 37°C 10 minutes. Samples were then digested by proteinase K at 70°C for 10 minutes and DNA was extracted by ethanol precipitation. Libraries were prepared using KAPA Hyper Prep Kit (KAPA) and custom Y-shaped TruSeq adapters according to the manufacturer’s instructions. All libraries were sequenced on a NextSeq 500 platform. Protein A-MNase (batch 6) and Yeast spike-in DNA were kindly provided by Dr. Steve Henikoff. The antibodies used were anti-NFIA (Deneen laboratory, (Fancy et al., 2012)), anti-NFIA (Sigma Aldrich, HPA006-111), H3K27me3 (Cell Signaling, 9733S), H3K27ac (Abcam, ab4729), and anti-Nkx6-1 (DSHB, F55A10).

### CUT&RUN data analysis

Raw paired-end reads were aligned to the mm9 genome according to (Skene et al., 2018). Briefly, fastq files were mapped using Bowtie2 (version v2.2.5) with the following options: --local --very-sensitive-local --no-unal --no-mixed --no-discordant --phred33 -I 10 -X 700. Peaks were called from aligned BAM files and focused using HOMER findPeaks passing a poisson P-value threshold of 0.0001. Blacklisted mm9 regions were removed using bedtools subtract before annotating peaks using HOMER annotatePeaks. For the final NFIA CUT&RUN peak calling, mm9 aligned BAM files derived from both NFIA antibody pulldowns were cross-referenced.

### Rescue experiment

Rab39b cDNA was cloned using Gibson Assembly into pEGF-N1-Flag plasmid (Addgene #60360, Britton et al. Nucleic Acid Research 2014). Pancreatic islets were isolated from age- and gender-match NPKO and controls mice using collagenase perfusion. Twenty islets were dissociated and plated on one-well od 804G collated 96-well plates for overnight recovery in RMPI media supplemented with 10% FBS. Next day cells were transfected using Lipofectamine 2000 and 100 ng of Rab39-or empty backbone-plasmid DNA per well. Cells were maintained for 6 days and then subjected to the GSIS. Insulin secreted in media was measured by ELISA (Alpco) and normalized to cell number and basal insulin secretion (after 2.8 mM glucose challenge).

### Statistical analysis

P values were calculated as indicated in figure legends using two-sided Student t-test. Data are presented as mean ± SEM and the following symbols are used to represent p values, \**p*<0.05, \*\**p*< 0.01, \*\*\**p*<0.005, and *\*\*\*\*p*<0.001. N represents number of independent experiments.

