## Supplementary figures and images for "NFIA regulates granule recruitment and exocytosis in the adult pancreas"

### Figure S1

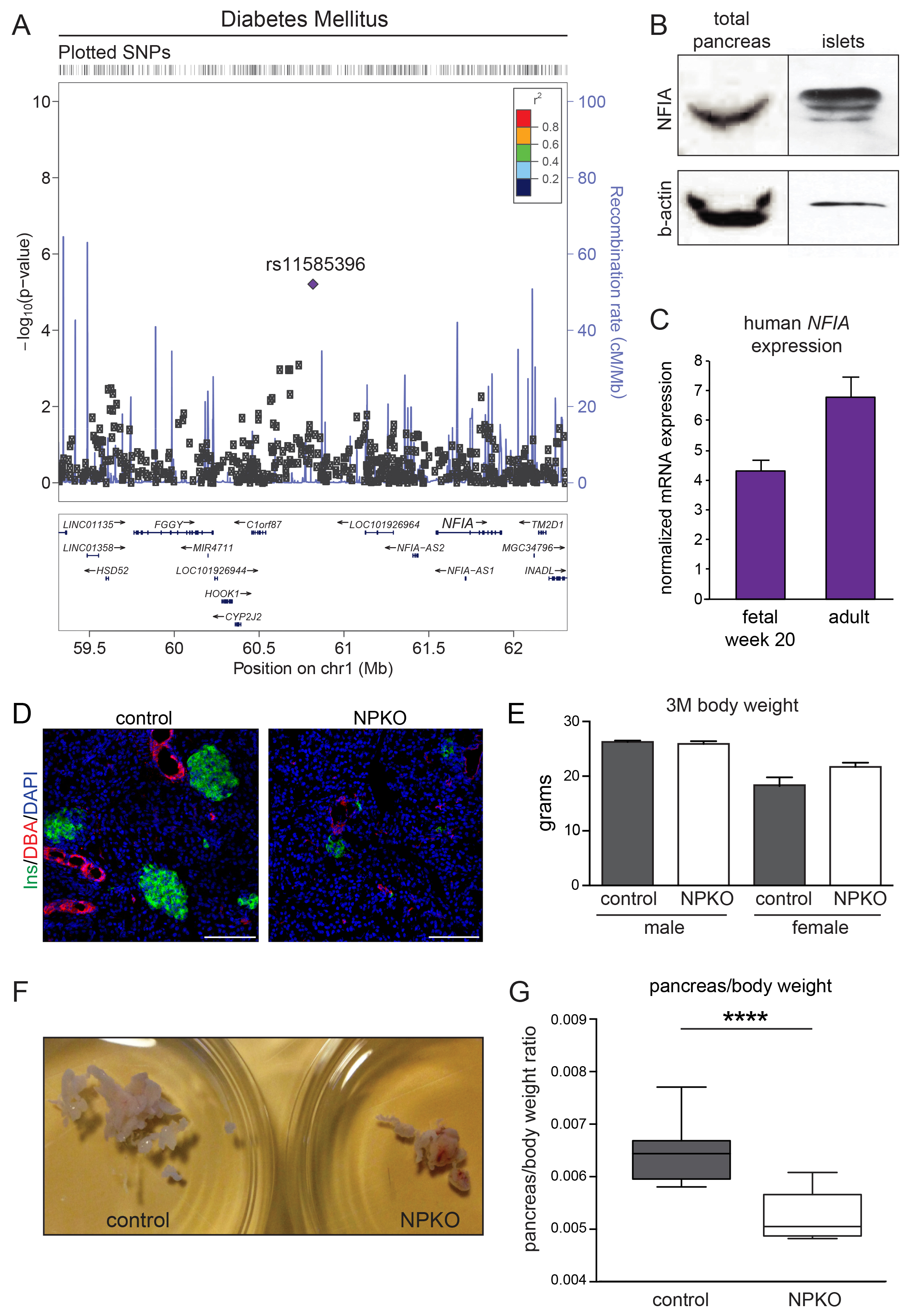

### Figure S2

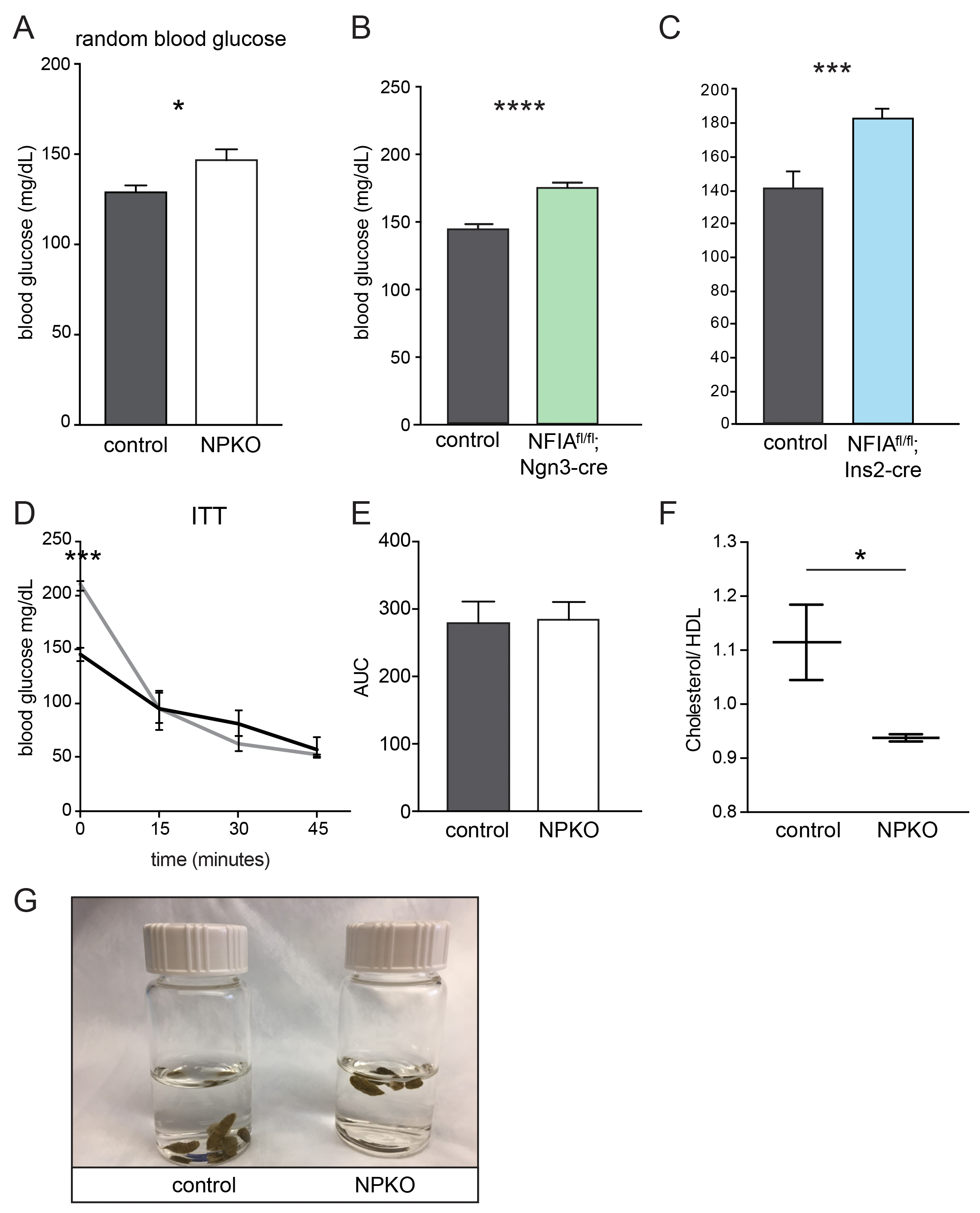

### Figure S3

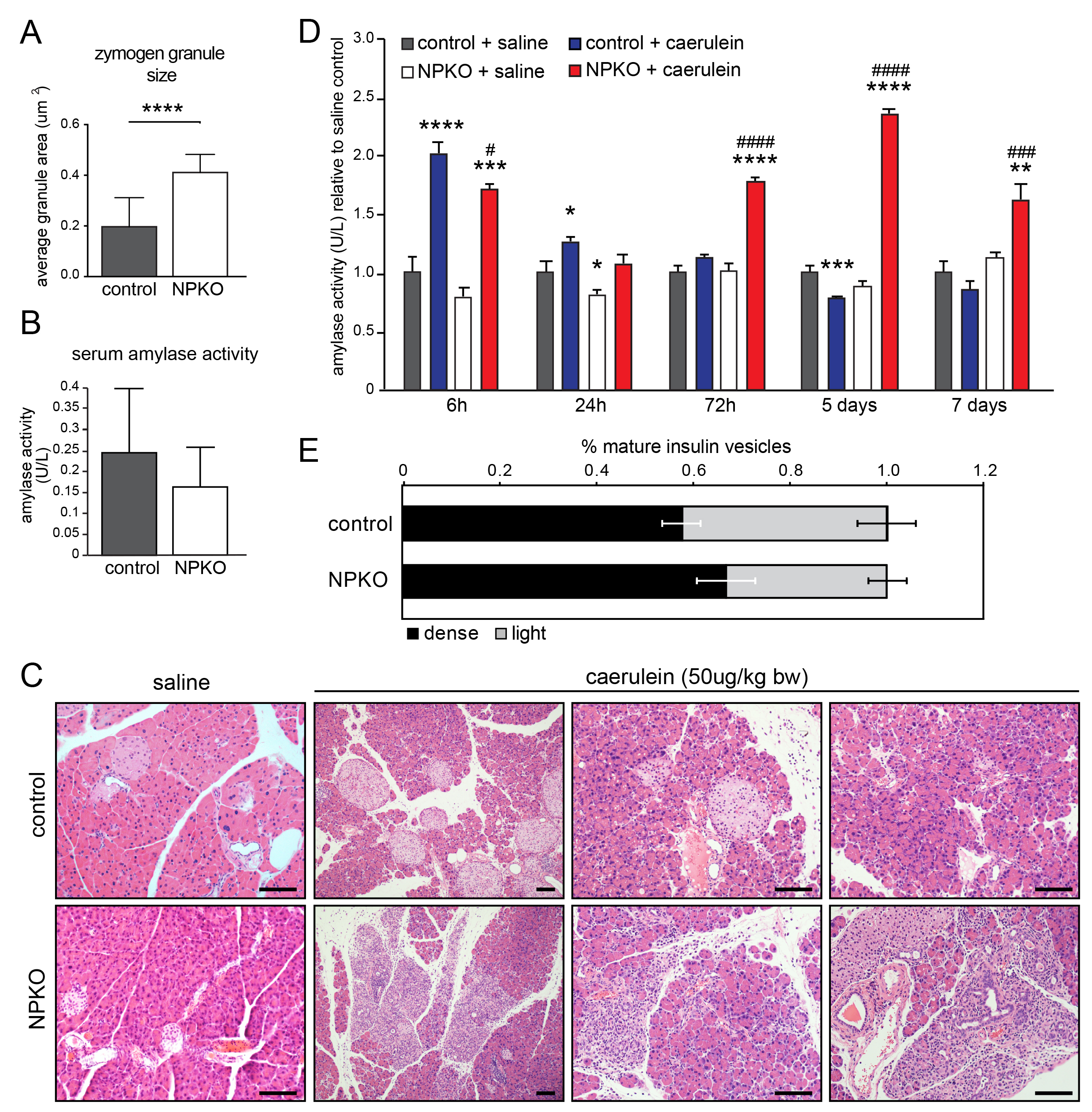

### Figure S4

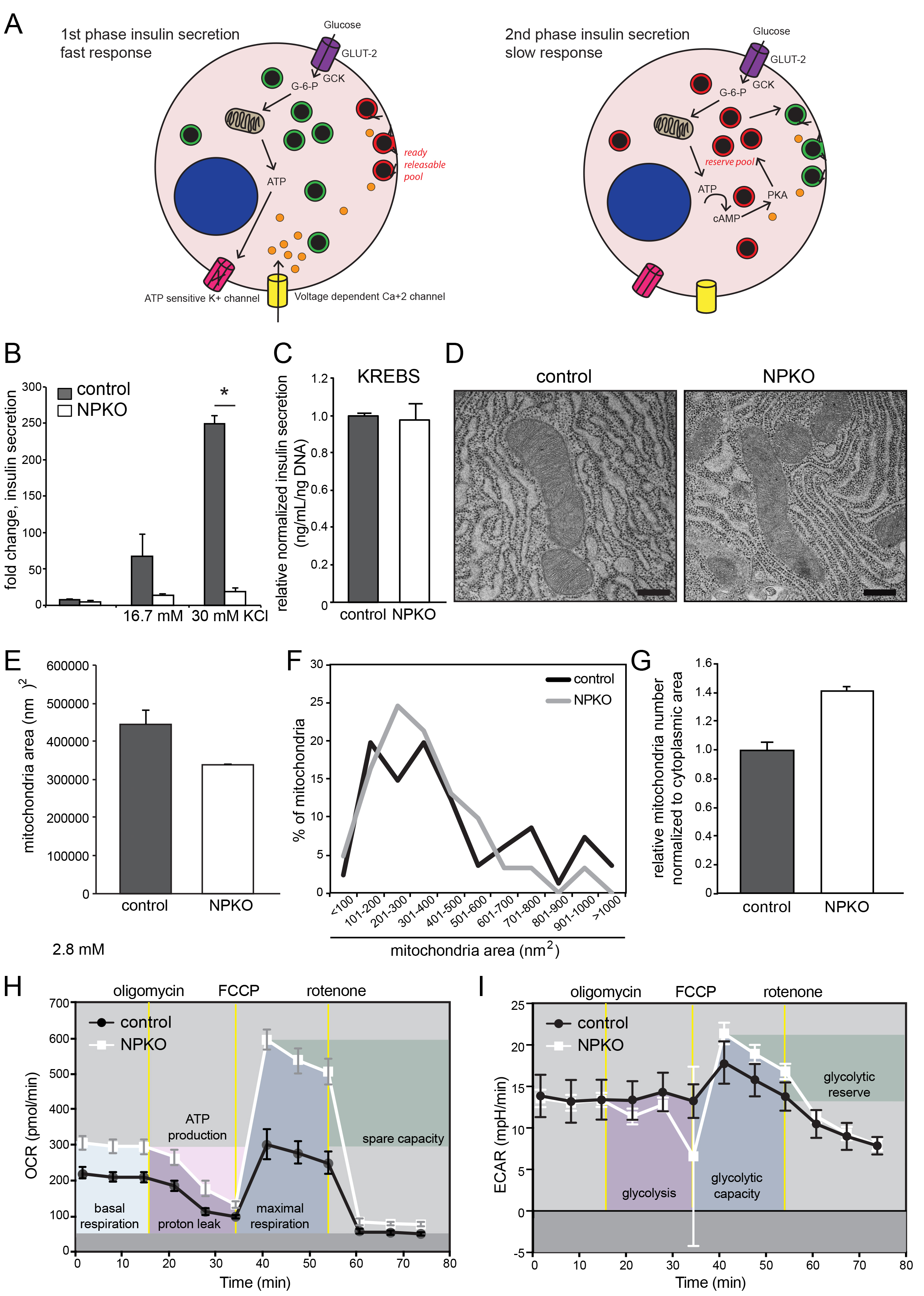

### Figure S5

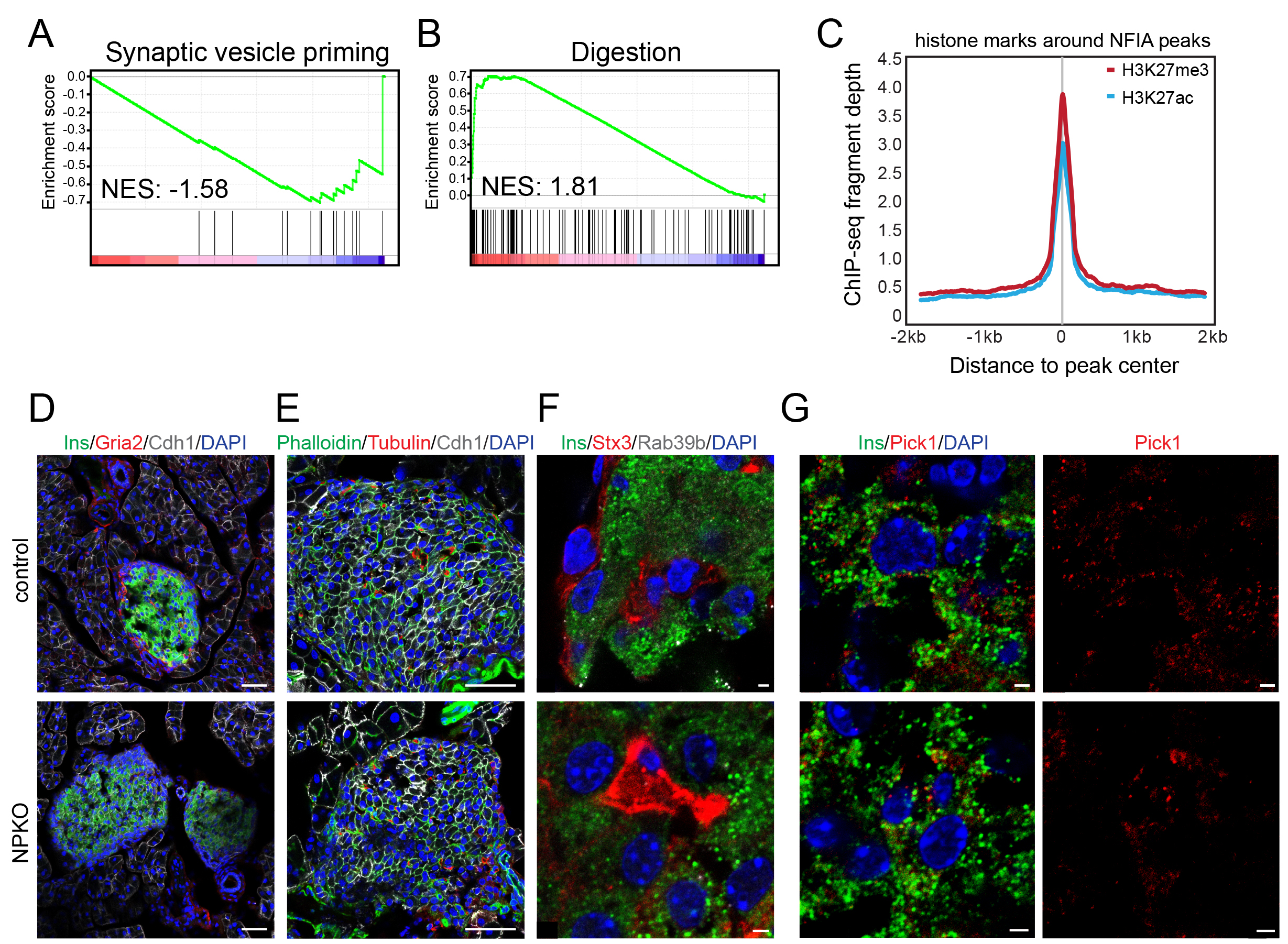
